# Get rid of the beat in mobile EEG applications: A framework towards automated cardiogenic artifact detection and removal in single-channel EEG

**DOI:** 10.1101/2021.02.09.430184

**Authors:** Neng-Tai Chiu, Stephanie Huwiler, M. Laura Ferster, Walter Karlen, Hau-Tieng Wu, Caroline Lustenberger

**Affiliations:** School of Medicine, National Yang Ming Chiao Tung University, Taiwan; Neural Control of Movement Lab, Department of Health Sciences and Technology, ETH Zurich, Zurich, Switzerland; Mobile Health Systems Lab, Department of Health Sciences and Technology, ETH Zurich, Zurich, Switzerland; Department of Mathematics, Duke University, Durham, NC 27708, USA; Department of Statistical Science, Duke University, Durham, NC 27708, USA; Mathematics Division, National Center for Theoretical Sciences, Taipei, Taiwan

**Keywords:** Electroencephalogram, Cardiogenic artifact, automated artifact removal

## Abstract

Brain activity recordings outside clinical or laboratory settings using mobile EEG systems have recently gained popular interest allowing for realistic long-term monitoring and eventually leading to identification of possible biomarkers for diseases. The less obtrusive, minimized systems (e.g. single-channel EEG, no ECG reference) have the drawback of artifact contamination with varying intensity that are particularly difficult to identify and remove. We developed brMEGA, the first algorithm for automated detection and removal of cardiogenic artifacts using non-linear time-frequency analysis and machine learning to (1) detect whether and where cardiogenic artifacts exist, and (2) remove those artifacts. We compare our algorithm against visual artifact identification and a previously established approach and validate it in one real and semi-real datasets. We demonstrated that brMEGA successfully identifies and substantially removes cardiogenic artifacts in single-channel EEG recordings. Moreover, recovery of cardiogenic artifacts gives the opportunity for future extraction of heart rate features without ECG measurement.

## Introduction

Wearable and portable EEG technology emerged in the last decades allowing for brain activity monitoring in a natural setting. Moving from clinical and laboratory settings into in-home settings opens up a vast amount of opportunities for real-life, long-term and large-scale monitoring of brain activity during different vigilance states and conditions, such as during sleep or while performing specific everyday tasks. Ultimately, this technology has the potential to serve as a digital biomarker for specific disorders, to detect in real-time aberrant brain activity (e.g. epileptic seizures), to better understand the functional role of brain oscillations in behavior and restoration, and to enable brain-computer interfaces in-home settings (e.g. rehabilitation, neurofeedback^1–4^).

In order to integrate this technology in real-life settings, the developed systems thrive for reduced obtrusiveness. This is achieved with miniaturization, reduction of EEG electrodes to a minimal number (sometimes only containing one EEG derivation), and localization of the electrodes to easy reachable places on the head (e.g. electrodes on forehead, mastoid, around-ear, and in-ear^5–7^). While reduction of electrodes increases comfort to the user it also comes with different drawbacks. One of them is the increased difficulty to separate real brain signal from noise and artifacts. EEG can be strongly contaminated by different artifacts, either from physiological (e.g. cardiogenic, sweating) and non-physiological sources (e.g., power-line noise)^8^. Especially when the artifacts have components in the brain activity frequency ranges of interest, they need to be removed for an accurate estimation of underlying brain activity.

The cardiogenic artifact in EEG can occur when cardiac electrical fields affect the surface potentials on the scalp^9^. It can arise in all vigilance states, e.g. also during sleep that is otherwise less affected by movement artifacts or eye blinks. The cardiogenic artifact has frequency components in the delta-theta range but due to its non-sinusoidal shape, the artifact is also represented in higher harmonics of the initial band leading to artifacts across the whole power spectrum. The affected frequency bands are of fundamental interest in neuroscientific research (e.g. slow-frequency activity in sleep, theta activity in wakefulness). Furthermore, cardiogenic artifacts are typically observable as a QRS shaped form, yet they can strongly vary in size and form. This adds an additional layer of difficulty in successfully detecting and removing them^9^.

Removal of cardiogenic artifacts necessitates (1) determination of existence of cardiogenic contamination in the EEG; (2) when cardiogenic artifacts exist, determine their locations; (3) removal of the artifact from the EEG (including correctly identifying cardiogenic artifact locations and separating the artifact from the real brain activity). For (1), to our knowledge, there is very limited work aiming to automatically determine whether cardiogenic artifacts exist when only a single-channel EEG signal is available. Tamburro et al. (2019) and Shao et al. (2009) proposed a framework for multi-channel cardiac artifact detection and removal using independent component analysis (ICA)^10,11^. Other approaches include hybrid blind-source separation and machine learning algorithm^12^, polynomial neural network and decision tree^13^, and feature selection-based approach^14^. For all these listed approaches multiple EEG channels are also required and are therefore not suitable for single-channel EEG applications. For (2), there have been some works with only single-channel EEG applications^15,16^. When the cardiogenic artifacts are *known to exist*, different approaches have been used to define the presence of the QRS complex without ECG reference including the energy interval histogram^15^, a modified S-transform approach^16^, and the continuous wavelet transform algorithm (CWT)^17^. Yet, all the approaches did not provide an application to tackle (1) and only well-controlled recording segments with known ECG artifact presence were included. We emphasize that while (1) seems to ask the same problem as (2), they are essentially different. For (3), several methods have been proposed to remove the cardiogenic artifact in EEG specifically when there is more than one EEG channel, or when at least an extra ECG reference channel is available^18–21^. To date, there is limited work when only a single EEG channel was recorded. The proposed methods to remove cardiogenic artifacts in single-channel EEG could be classified into two categories. The first category includes algorithms related to the template subtraction (TS), or ensemble average subtraction^15,16,22^ when the locations of cardiogenic artifacts are known; for example, the modified template subtraction by weighting^15^ or shape adaptive nonlocal artifact removal (SANAR)^23^. The second category includes algorithms generalizing ICA to the single-channel setup; for example, the single-channel ICA (scICA)^24^, the wavelet ICA (wavelet-ICA)^25^, and the ensemble empirical mode decomposition ICA (EEMD-ICA) is combined in Mijovic et al. (2010)^26^. While these ICA based algorithms might work, they are limited by the need of visual inspection to select channels for the cardiogenic artifact removal and sometimes machine learning tools are considered to automatize the mission of artifact component selection. Also, it is not easy to avoid the mode mixing issue when the wavelet or EEMD decomposition is applied. Moreover, the lack of theoretical justification of EEMD further limits its application for scientific research.

Correct removal of artifacts initially requires identification of participants and epochs that are contaminated by EEG. Even though similar EEG montages might be used, not all participants will show artifacts. Furthermore, specifically for long-lasting recordings (e.g. nocturnal sleep period), these artifacts might only temporarily be present and will presumably vary in their magnitude and shape across the recording period. Specifically, for large-scale and long-term recordings that are collected with mobile applications, visual inspection (which is typically involved) is not feasible.

Collectively, there is still lack for a framework that involves automated identification and removal of cardiogenic artifacts from single-channel EEG that have been collected over longer recording durations and without an ECG reference signal. Yet, such a framework is urgently needed, given that mobile, non-obtrusive EEG applications have gained tremendous interest in research. In order to fill this gap, we combine here for the first time nonlinear-type time-frequency analysis tools and machine learning to automatically determine whether a cardiogenic artifact exists and to remove it from single-channel EEG without the need of an ECG reference. We first train a machine learning algorithm based on novel features extracted from nonlinear-type time-frequency analysis tools to determine whether cardiogenic artifacts exist. Second, we apply a manifold learning algorithm similar to SANAR^23^ to remove cardiogenic artifacts. The algorithm is modified to handle the specific characteristics of cardiogenic artifact--unlike the stimulation artifact removal mission, the locations of cardiogenic artifacts are in general unknown. We coin our algorithm ***b****eat* ***r****emoval* ***m****obile* ***E****E****G a****lgorithm* (brMEGA). We apply brMEGA to long-term, nocturnal EEG recordings that were obtained from a portable device in a participant administered in-home study and validate our framework in datasets for which ECG reference channels exist. We demonstrate that contaminated EEG can be successfully identified and substantially restored with our proposed framework.

## Results

Here we included thirteen nocturnal single-channel EEG recordings that showed various degrees of cardiogenic artifact contamination, both across and within recordings and that were obtained from a mobile EEG device, the MHSL-Sleepband^6^. In addition, two additional datasets with and without cardiogenic contamination from a sleep lab were included that also had synchronous ECG channels from the whole night polysomnogram. A summary of the proposed cardiogenic artifact removal algorithm, brMEGA, is provided in Figures 1 and 2. Details about the included dataset and established algorithms can be found in the Methods section. The MATLAB source code used in this publication is available via request.

**Figure 1:**
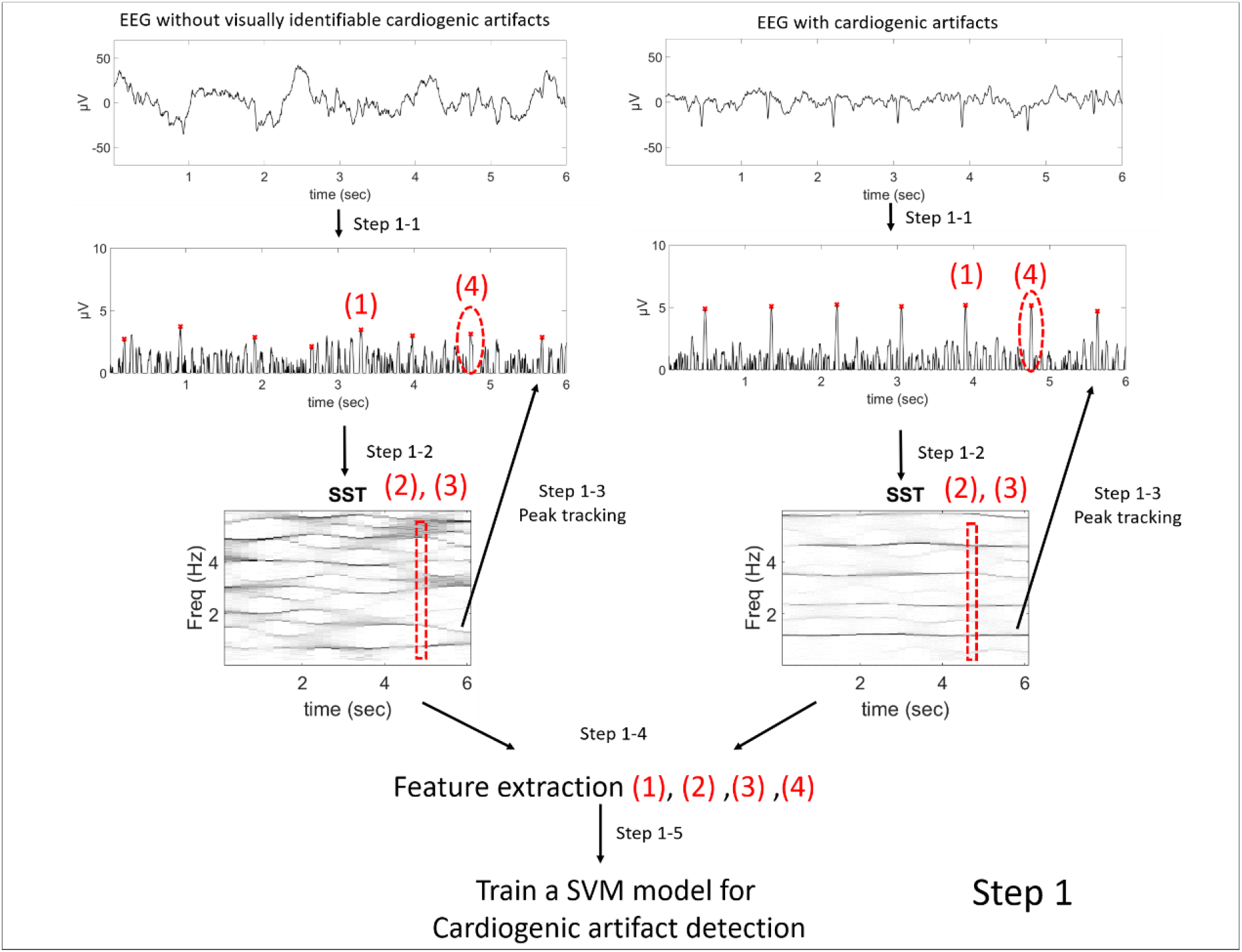
The flowchart of the first step of brMEGA. The four designed features are marked from 1 to 4. The first feature is the peak height in the time domain, the second and third features are the features in the time-frequency (TF) domain that are determined by the “optimal transport distance”, and the fourth feature is the total energy around the peak. The features in the time domain depend on the peak tracking algorithm with the extracted instantaneous frequency from the TF representation.

**Figure 2:**
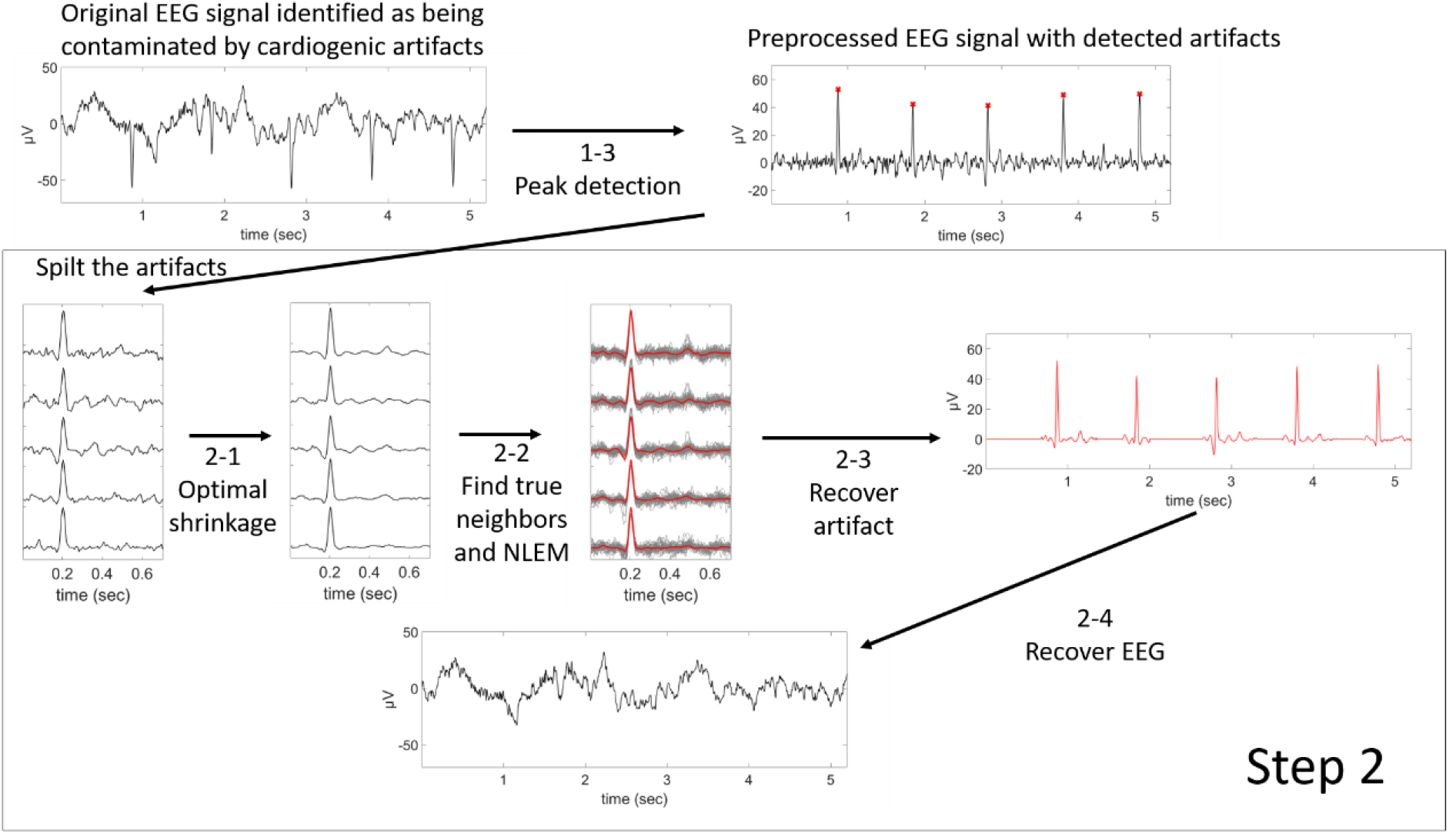
The flowchart of the second step of brMEGA.

### Non-linear time-frequency analysis and machine learning enable automatic determination of existence of cardiogenic artifacts in EEG (Step 1)

In the first step of brMEGA, we aimed at automatically determining if cardiogenic artifacts exist. Particularly, without the ECG signal or cardiac rhythmic information, we need to estimate where the cardiogenic artifacts are. However, due to the variability of the heart rate, the traditional time-domain and spectral approaches are limited since the cardiac rhythm is not fixed. To handle these challenges, we take a well-known fact into account. When cardiogenic artifact exits as a periodic component, the time-frequency representation (TFR) of the EEG signal with cardiogenic artifact contains dominant curves associated with the cardiogenic artifacts; otherwise, there is no dominant curve. This specific characteristic in the TFR is converted into features so that we can learn if cardiogenic artifacts exist. To do so, the signal is composed with the rectified linear unit activation function and then smoothened (step 1-1). Then we apply the recently established non-linear time-frequency methods of synchrosqueezing (SST)^27–29^ (step 1-2) to extract dominant spectral features from the TFR, particularly the instantaneous frequency (IF). When the cardiogenic artifacts exist, the IF is the instantaneous heart rate (IHR), so we obtain the IHR from SST at the same time; otherwise, we obtain the IF associated with the EEG spectrum. With the IF, the peak tracking algorithm is applied to obtain the peak locations (step 1-3). When cardiogenic artifacts exist, we obtain the cardiogenic artifact locations; otherwise, we obtain random EEG spikes. From steps 1-2 and 1-3, we form four features associated with cardiogenic artifacts, two from the time domain and two from the TFR (step 1-4). We apply the well-established support vector machine algorithm (step 1-5) to learn whether a 4-second epoch is contaminated by cardiogenic artifact or not. The flowchart of this part of algorithm is shown in Figure 1.

The neuroscientist expert in our team visually examined the EEG signal and provided a quality index describing the strength of cardiogenic artifact over non-overlapping 4 seconds epochs. An epoch was labeled as 0 if no clear cardiogenic artifacts could be visually identified, and 1 if a cardiogenic artifact was clearly visible. For 13 whole night polysomnograms (all from different participants), the neuroscientist labeled randomly selected 100 4-second epochs between the first and second hour of each recording.

There were in total 1300 labeled 4-second epochs from 13 subjects. The leave-one-subject-out cross validation result is shown in Table 1, where the sensitivity is 0.95, the specificity 0.70, and the F1 score is 0.84 and the overall accuracy is 0.81. The 5-fold cross validation carried out over all epochs from all subjects revealed similar performance values (Appendix Table A1).

**Table 1:**
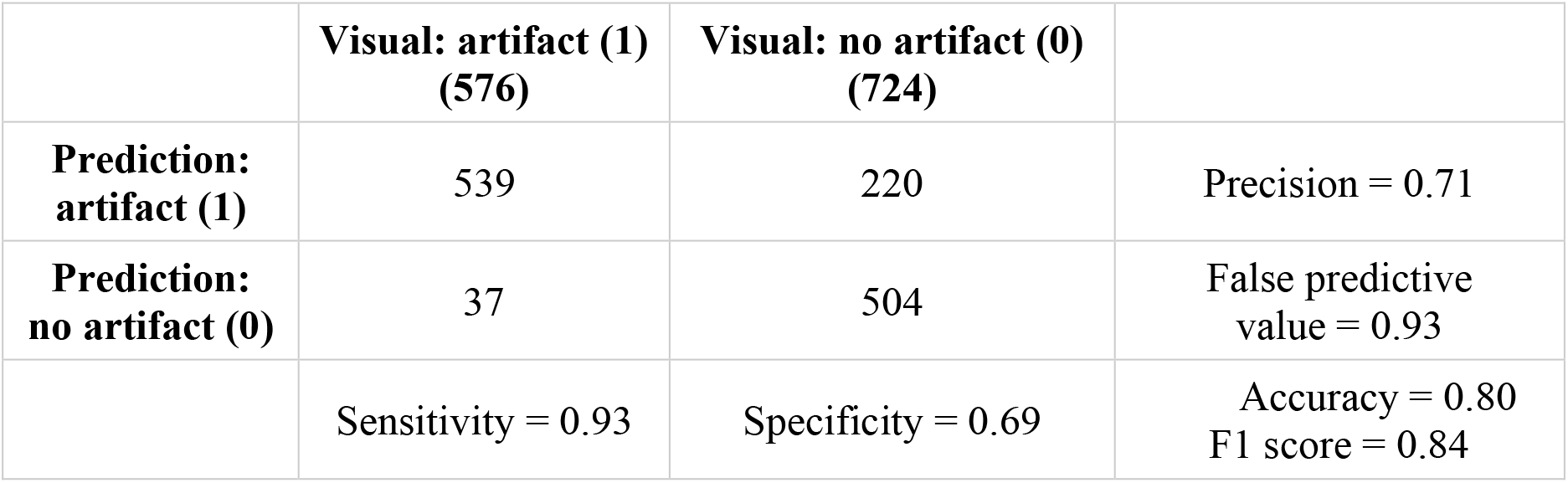
The overall confusion matrix of the leave-one-subject-out cross validation.

|  | **Visual: artifact (1)**<br>**(576)** | **Visual: no artifact (0)**<br>**(724)** |  |
| --- | --- | --- | --- |
| **Prediction:**<br>**artifact (1)** | 539 | 220 | Precision = 0.71 |
| **Prediction:**<br>**no artifact (0)** | 37 | 504 | False predictive<br>value = 0.93 |
|  | Sensitivity = 0.93 | Specificity = 0.69 | Accuracy = 0.80<br>F1 score = 0.84 |

### Template subtraction (TS) using the wave-shape manifold model can accurately remove cardiogenic artifacts of different sizes and forms (Step 2)

In the second step of brMEGA, we aimed at removing the cardiogenic artifacts. Once an epoch is determined to be contaminated by cardiogenic artifacts in step 1, we apply the beat tracking algorithm combined with the estimated IHR by SST to obtain the location of cardiogenic artifacts (step 1-3). With these locations, we move forward to remove the cardiogenic artifacts. A first glance at the cardiogenic artifact contaminated EEG signal suggests that the cardiogenic artifacts share a common spike-like pattern. This common pattern motivated the TS algorithm. The traditional TS approach, however, is limited when the spike-like patterns are not fixed from one to another, which is the case in the cardiogenic artifacts in EEG. To fix this issue, the manifold learning algorithm was introduced. As a generalization of linear model, a manifold model is capable of describing the nonlinear relationship among cardiogenic artifacts^30^. Based on this model, we apply the optimal shrinkage (OS) to find similar cardiogenic patterns (step 2-1), so that the nonlocal Euclidean median (step 2-2) leads to a less biased cardiogenic artifact estimates. Unlike the traditional TS approach that mainly counts on the previous few cardiogenic artifacts, this step allows us a better reconstruction of spike-like cardiogenic artifacts (step 2-3) and hence a more accurate EEG recovery (Step 2-4). The flowchart of this part of algorithm is shown in Figure 2.

Figure 3 illustrates the TF representation of one representative nocturnal recording before and after cardiogenic artifact removal. All other 12 nights can be found in Supplementary Figures A1-A12. Application of the brMEGA recovers the clean EEG by substantial removal of the artifacts in the delta-theta and high frequency region (including higher-harmonics). In the temporal domain, this finding is confirmed by an accurate recovery of the signal (see Figure 4 for an exemplary temporal trace).

**Figure 3:**
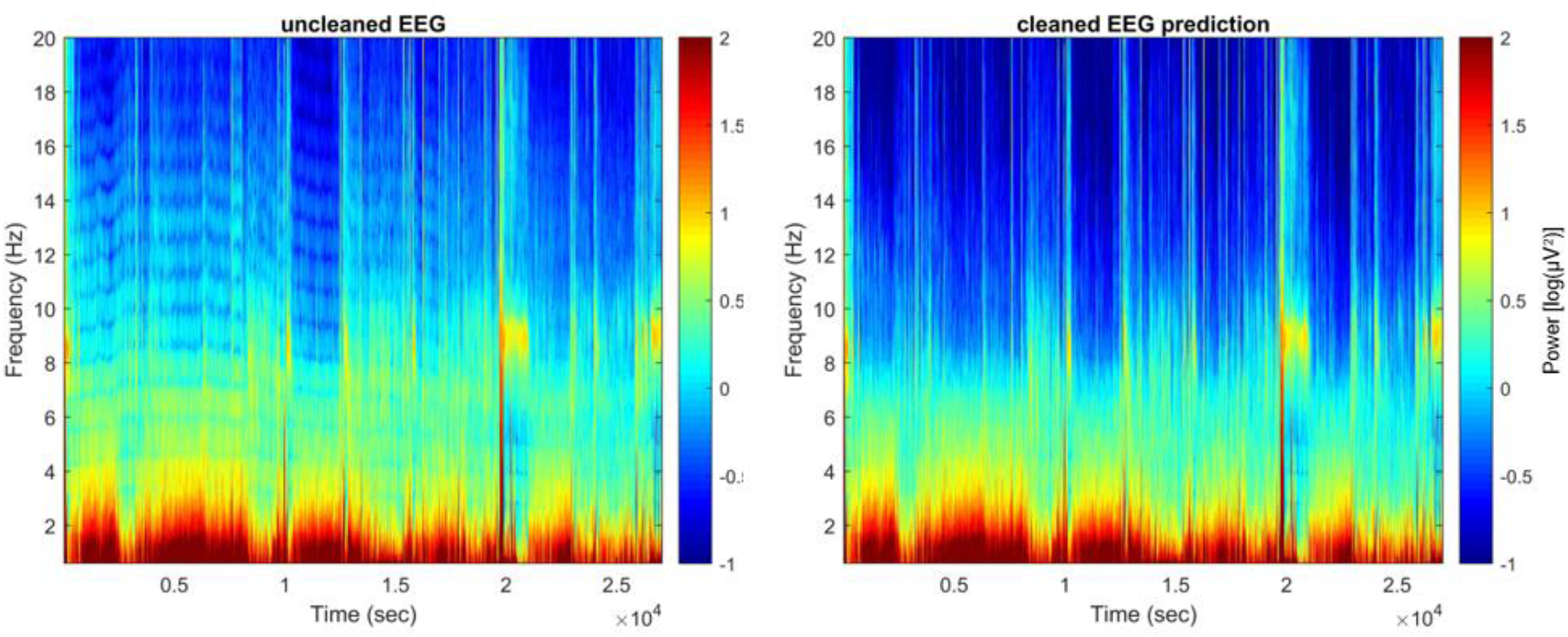
TF representation of the original EEG signal (all-night nocturnal recording, left) and after removing the cardiogenic artifact (right).

**Figure 4:**
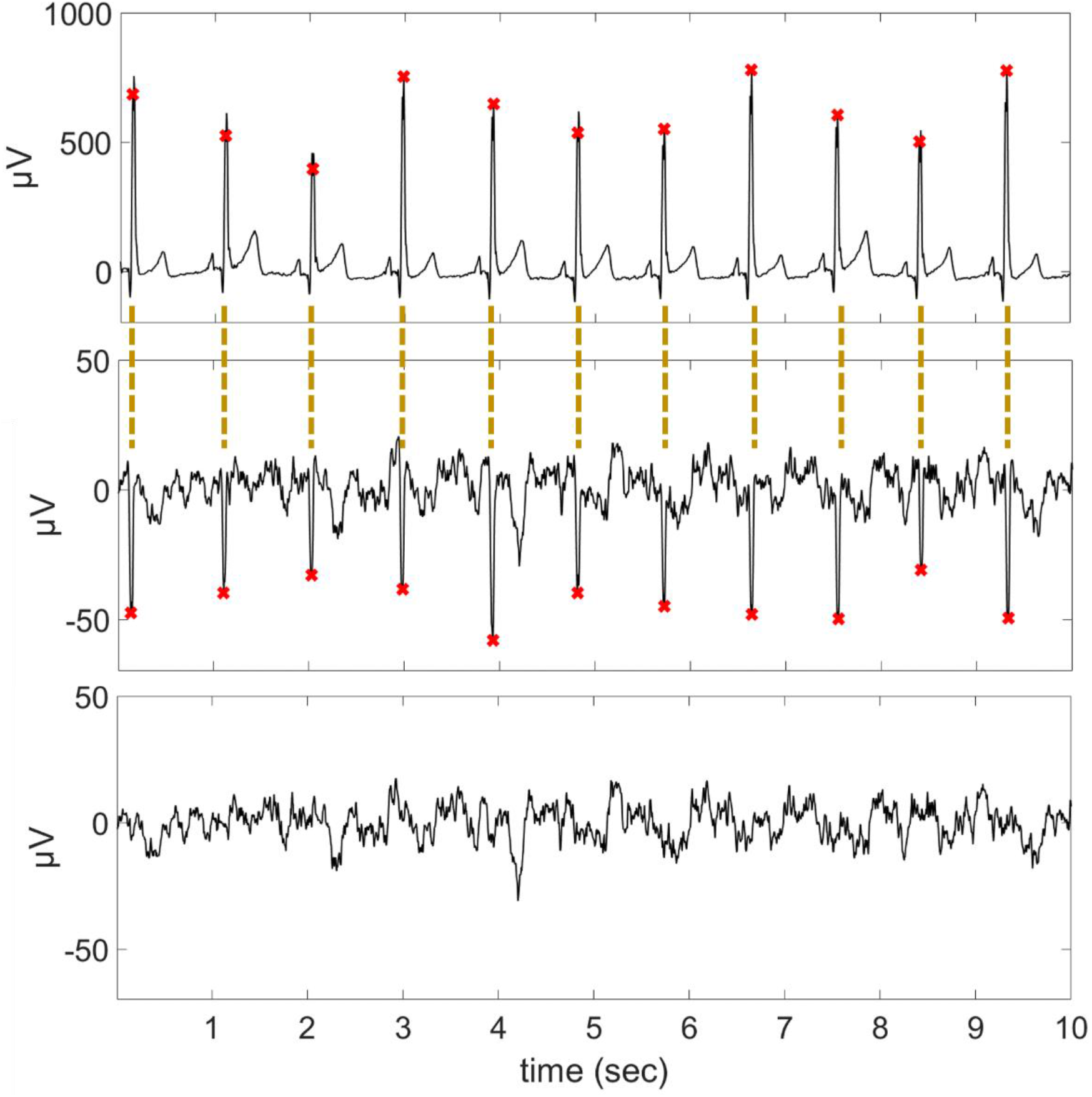
Illustrations of the cardiogenic-artifact contaminated EEG signal (second panel) and the simultaneously recorded ECG signal (top panel). Red x marks the respective peak detection for both ECG (R peak) and EEG (cardiogenic artifact peak) separately. The R peak and artifact peaks are correctly identified and overlap. The recovered EEG signal by brMEGA is shown in the third panel.

### ECG QRS location and structure match well with algorithm identified artifact

In order to validate the performance of our brMEGA to accurately define ECG peak locations we included two in-lab measured subjects from one study site that also included ECG as reference recordings. In Figure 3, a representative segment before and after cardiogenic artifact removal is shown, and the simultaneously recorded electrocardiogram is illustrated for a visual comparison. Visually identified cardiogenic artifacts coincide with QRS complexes in the ECG signal. To quantify the performance^31^, the R peaks of the ECG are viewed as the ground truth. To avoid the impact when the ECG signal is of low quality, the ECG signal quality is determined by bSQI^32^. We keep all epochs when the associated ECG signal is of bSQI>0.95 in the final evaluation, 4.4% epochs are removed. As is illustrated in Table 2, the detected cardiogenic artifact peaks in two cases by brMEGA correspond with 96% precision and 96% sensitivity to R peaks defined in the ECG with the grace period of 50 ms. The mean absolute error is 1.24 ms. We therefore conclude that brMEGA can detect the existence of cardiogenic artifact and remove cardiogenic artifacts from the single-channel EEG signal.

**Table 2.** The confusion matrix of cardiogenic artifact detection. A detected cardiogenic artifact is viewed as a false positive when there does not exist a R peak within the 50 ms grace period. A false negative cardiogenic artifact is defined when there is a R peak, but we do not detect a cardiogenic artifact within the 50 msec grace period.

|  | **ECG R peak**<br>**(number = 39,942)** |  |  |
| --- | --- | --- | --- |
| **Detected EEG**<br>**cardiogenic artifact**<br>**(number = 39,730)** | True positive<br>number = 38,262 | False positive<br>number = 1,468 | PPV = 0.96 |
|  | False negative<br>number = 1,680 |  | F1 = 0.96 |
|  | Sensitivity = 0.96 |  | MAE = 1.24 |

In Figure 5, an additional comparison of the recovered cardiogenic artifacts and ECG signal is demonstrated. It is noteworthy that in addition to the dominant spike associated with the R peak in the ECG signal, there is a bump after the dominant spike in the reconstructed cardiogenic artifact (indicated by red arrows). Based on the temporal characteristics of a classical ECG curve as derived from our averaged ECG signal, the second bump observed in the cardiogenic artifact is likely associated with the T wave in the ECG signal.

**Figure 5:**
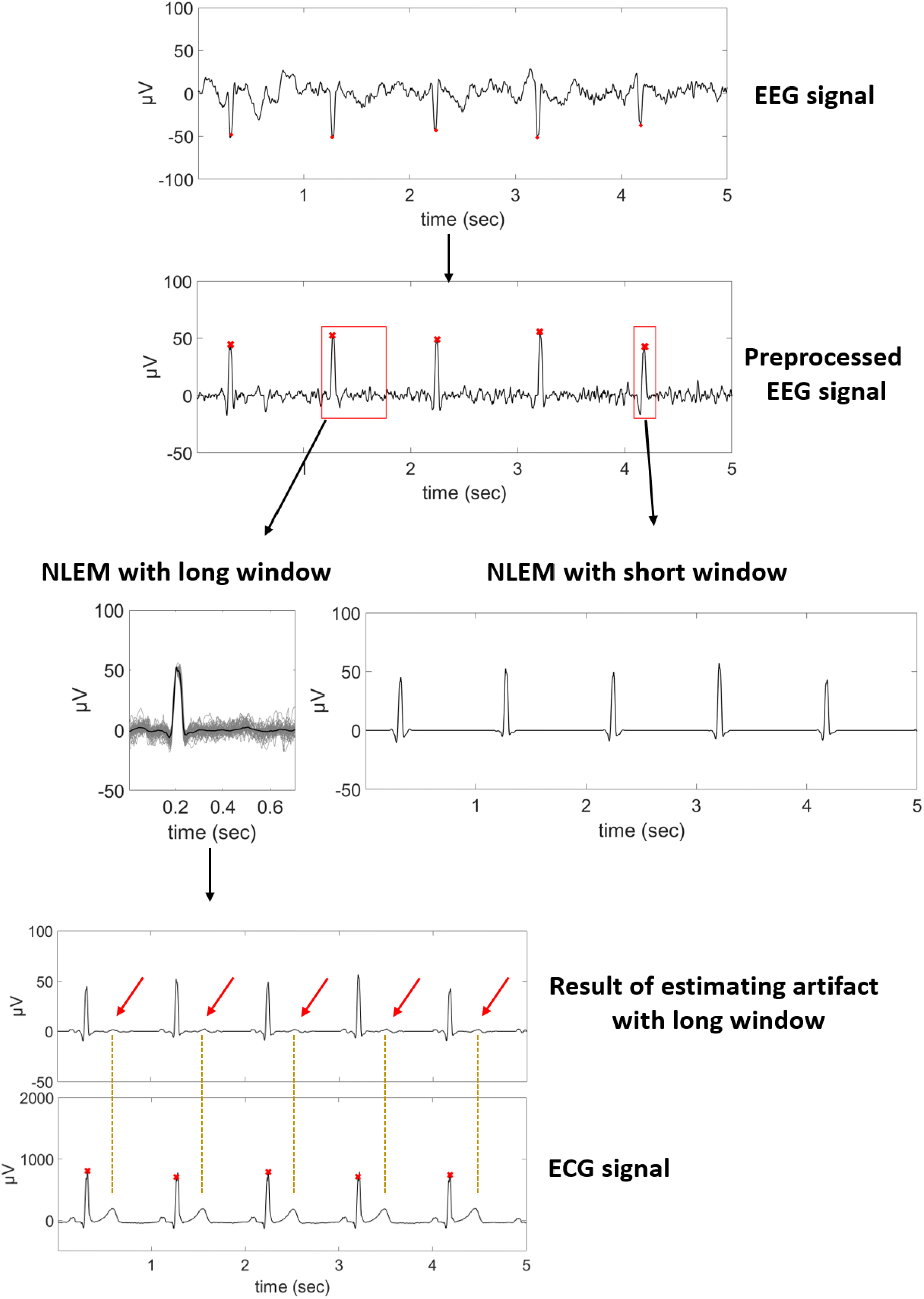
An illustration of the cardiogenic artifacts and the “T wave” bumps (marked by red arrows). If the window is short, the T-wave-like artifact cannot be recovered. Red x indicate cardiogenic artifact peak or R peak occurrence.

### Manifold TS outperforms previously published TS approach in removing the cardiogenic artifacts from contaminated EEG

In addition to our proposed brMEGA algorithm, we also show the result by a previously published TS approach^15^, which is called ensembled average subtraction (EAS). For a fair comparison, step 2 of the proposed brMEGA is replaced by EAS.

In order to quantify and compare the TS performance, we consider the following two indices: the *artifact residue (AR)* index, which was considered in Malik, et. al.^33^ for the f-wave extraction from ECG signals, and a modification of the *spectral concentration* (SC) index proposed in Castells, et. al.^34^ (details in methods section). The quantitative results shown in Figure 6 indicates that overall the proposed brMEGA algorithm outperforms the existing EAS algorithm as demonstrated by a significant reduction in both SC and AR (values get closer to 0) across all participants. Note that 2 out of 13 cases have less than 30 epochs contaminated by cardiogenic artifacts (S029: 24/1562 and S053: 2/1669), so they are not shown and summarized below. Overall, the mean and standard deviation of the AR index for brMEGA is 0.04±0.015, while those for EAS is 0.054±0.028. For each subject, the null-hypothesis that brMEGA does not perform better than EAS in the sense of AR index is rejected by the paired-sample t-test with the p value < 10^−10^. Regarding the SC index, the mean and standard deviation for brMEGA is 0.013±0.003, while those for EAS is 0.015±0.003. For each subject, the null-hypothesis that brMEGA does not perform better than EAS in the sense of SC is rejected by the paired-sample t-test with the p value < 10^−5^.

**Figure 6:**
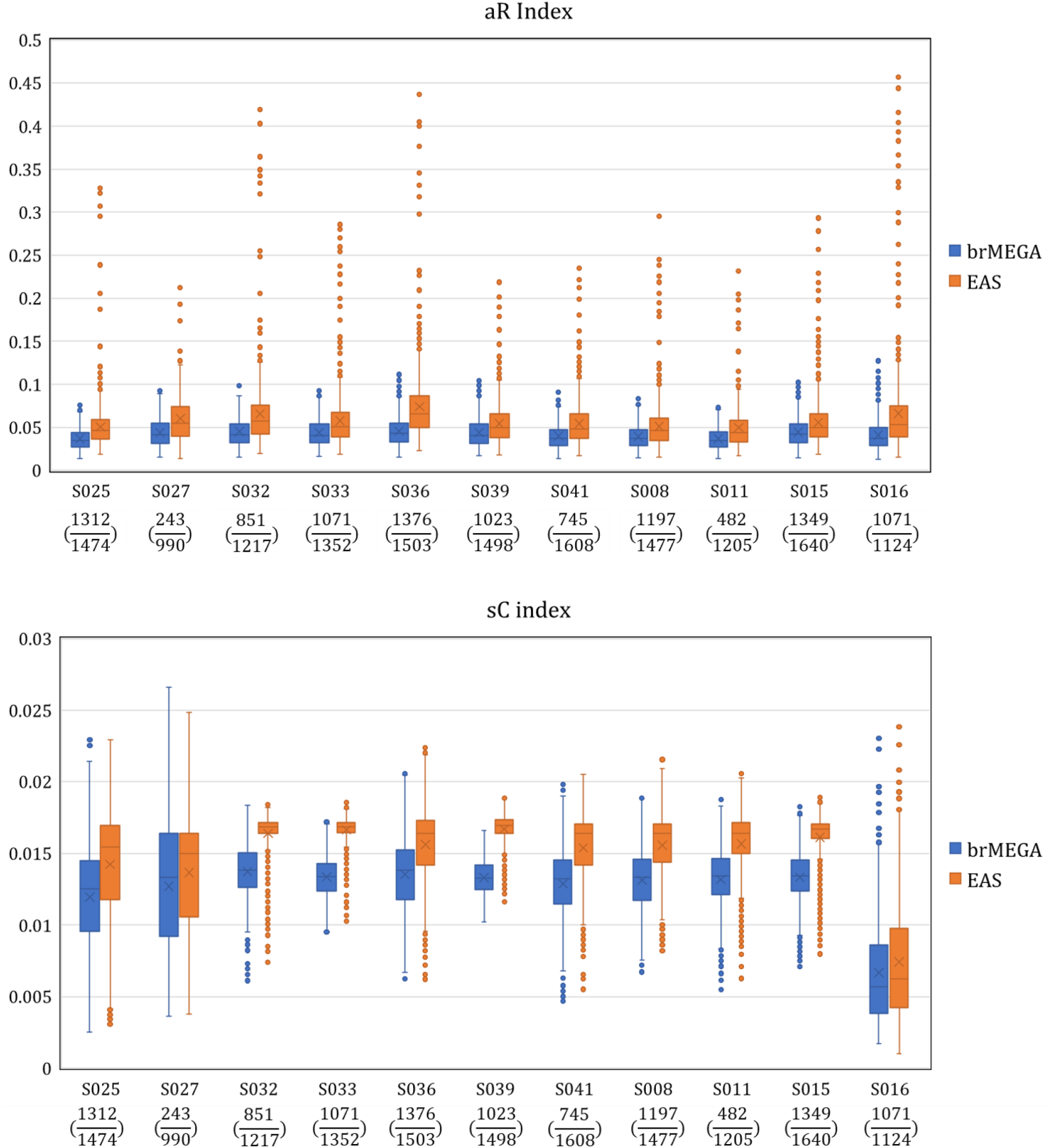
The AR and SC indices of 11 cases in which visually ECG artifact have been identified in the course of the nocturnal recording. The number of 20-sec epochs of each case identified as been contaminated by cardiogenic artifacts, and the total number of 20-sec epochs are listed below the case ID as the numerator and denominator respectively.

### Manifold TS outperforms previously published TS approach in recovering the EEG signal from ECG artifact

To further confirm that the EEG is properly recovered, we apply brMEGA and EAS to the semi-real database. In this dataset, we combine cardiogenic artifact-free EEG with varying cardiogenic artifacts that are characterized by the range and variability of the cardiogenic artifacts reported in the previous paragraphs (7 nocturnal EEG recordings from different participants). This approach allows us to compare the EEG with contamination and after applying artifact removal (brMEGA and EAS) to the ground truth EEG without ECG artifacts. To do so, we report here four indices that quantify EEG recovery. First, NMSE (normalized mean square error) is substantially lower after artifact removal with brMEGA (0.151±0.113), and EAS (0.184±0.129) compared to the contaminated EEG (1.241±0.941). Second, the cross-correlation (p) between the ground truth EEG and the brMEGA cleaned EEG (0.926±0.058) and EAS cleaned EEG (0.916±0.06) strongly increases compared to the correlation with the cardiogenic contaminated EEG (0.69±0.118). Third, the signal-to-noise ratio (SNR) substantially increases after artifact removal with brMEGA (8.919±2.361) and EAS (8.130±2.536) compared to the contaminated EEG (−0.025±2.739). The same holds true for the fourth index, peak SNR (PSNR), with brMEGA (17.806±2.245) and EAS (17.041±2.416) cleaned EEG being substantially higher than the uncleaned EEG (8.888±2.651). Overall, both brMEGA and EAS substantially recover the EEG as indicated by all four indices being pronouncedly improved compared to the contaminated EEG. For each index, brMEGA-cleaned EEG significantly differs from the contaminated EEG in all four indices when performing a linear mixed effects model with p-values < 0.001. Furthermore, comparing the differences to the uncleaned signal, brMEGA outperforms the EAS approach for all four indices with p-values significantly for SNR, and pSNR p-values < 0.02, and trend-level for NMSE, p, p-values = 0.065 and 0.057, respectively.

## Discussion

Here we propose the automated framework brMEGA for determination, peak location identification, and removal of cardiogenic artifacts from single-channel EEG. We demonstrate that our algorithm can successfully detect and substantially recover EEG periods with cardiogenic artifacts without the need of an additional ECG derivation. BrMEGA is the first algorithm for automated cardiogenic artifact determination and removal that can be applied to long duration recordings with minimal EEG channel settings. Therefore, it provides a highly attractive artifact removal solution for mobile EEG applications that have gained tremendous impetus in the last decade.

### Automatic determination of existence of the cardiogenic artifact

In the first step, brMEGA determines a possible presence of a cardiogenic artifact in the EEG. The cardiogenic artifact varies in amplitude from being completely absent to strongly contaminating the EEG, both across and within different recordings. Classical spectral analysis is generally limited when the signals are with time-varying frequency and amplitude that need identification with high temporal and frequency resolution (e.g. IHR) such as is the case for cardiogenic artifacts. Therefore, a method is needed that overcomes the limitations of classical spectral analysis. We have recently established that SST can successfully isolate and characterize this kind of signal, even when the signal oscillates in a non-sinusoidal way (e.g. non-sinusoidal signals such as ECG)^28^. Accordingly, the brMEGA detection algorithm included a combination of features extracted from the raw signal and the SST to capture typical cardiogenic signatures. With these features we have trained an SVM to identify cardiogenic artifacts in EEG based on visual expert labelling. Compared to a neuroscience expert, the algorithm achieved a very high specificity (at ~95%) in identifying an artifact that was also identified visually. The sensitivity was slightly lower at around 70%, thus the algorithm identified some epochs as including cardiogenic artifacts in the EEG that were not labelled by the expert. However, this might be biased to the visual capacity of the expert. Artifacts could be less pronouncedly visible, specifically if strong EEG activity is present, even though they exist in the EEG. We took a closer look at some epochs that were only identified by the algorithm and could also see indices of cardiogenic artifacts that were labelled by the algorithm indication that at least in some epochs, the algorithm might outperform the scorer (see Appendix Figure A13). A limitation of the datasets derived from the mobile EEG device is that we did not have the ground truth ECG to evaluate whether the located cardiogenic peaks with our algorithm matched with real R-peaks in the ECG. To overcome this limitation, we include additional in-lab datasets that provided simultaneous EEG and ECG recordings. Applying our brMEGA algorithm to these recordings demonstrated highly precise peak overlap between the cardiogenic artifact and the ECG with 96% precision and 96% sensitivity. This result is comparable with the accuracy of previously published approaches, for which accuracies around 97-99% were reported^10,16^. Here, we have to consider the achieved performance is slightly lower than in previous publications because we include a whole night recording with some time periods that might not have had an artifact. In other words, we include step (1) in this metric and likely have some inaccuracy in the determination of 4s epochs with artifact contamination whereas the previous publications only report accuracy from segments where a clear visual artifact was present^10^, or for which cardiogenic artifacts were added to clean EEG^16^.

To date, only few previous studies introduced a framework to automatically determine the existence of and remove cardiogenic artifacts in recorded data with unknown/different levels of contamination and without ECG reference^10–12^. They applied ICA in high-density EEG and developed an algorithm that could define IC with ECG artifact. However, their approach cannot be applied in single-channel EEG settings that are common in mobile applications. ICA is computationally very intensive and not ideal for long-term recordings, (a) because very long recordings would take up too much computational power, and (b) ICA requires a certain level of signal stationarity that is not provided for hour-long sleep and wake recordings therefore requiring sophisticated windowing approaches. Our presented framework here overcomes these limitations and enables accurate definition of contaminated segments and peak location independent of recording length also for single-channel EEG.

### Removal and recovery of the cardiogenic artifact

Upon detection of cardiogenic artifacts, we applied the wave-shape manifold model template subtraction to remove the cardiogenic artifact. Our approach can substantially recover the EEG as illustrated in the time-domain but also in the frequency domain, for which we should removal in the primary frequency bands but also the affected higher harmonics. This is also quantified in four indices that depict EEG recovery in contrast to the ground truth EEG in our semi-real dataset. All indices show pronounced improvement after brMEGA application compared to the contaminated EEG. The proposed brMEGA is suitable to handle non-stationary time series, particularly when the signal is recorded for hours or longer. It is well known that the EEG signal is non-stationary. The cardiogenic artifact, on the other hand, is also non-stationary, due to the inevitable heart rate variability and the fact that the cardiogenic artifact strengths might vary from one to another. Furthermore, changes in head positions can affect the extent of the artifact, which is often happening during sleep or recordings that involve head movements^9^. Unlike the traditional TS algorithms, like EAS we compared in this work^15^ and its variations^16,22^, we leverage this non-stationarity to more efficiently remove cardiogenic artifacts. Such non-stationarity is captured by the wave-shape manifold, which quantifies how cardiogenic artifacts change from one cycle to another. Specifically, for each given cardiogenic artifact, we design a specific filter, the optimal shrinkage, to “clean up the EEG signal” so that the median is taken only over those cardiogenic artifacts that are similar to the given one. Note that unlike the traditional TS algorithms, those chosen cardiogenic artifacts for the median operator might be temporally far away. When there are multiple channels, the proposed brMEGA can be generalized. A further exploration of such generalization and its comparison with ICA will be reported in our future work.

Our modified TS outperformed the EAS reported in Park et al. as quantified by the AR and mSC index^15^. The first term of the AR index measures how accurate the EEG signal is recovered and we need to avoid over-smoothing. The second term of the AR index, on the other hand, measures the residue of the cardiogenic artifact. The product captures simultaneously how well the cardiogenic artifact is removed and how well the EEG signal is recovered. Compared with its application to iEEG stimulation artifact removal^23^ and f-wave extraction^33^, its application to our problem is natural without too much change. The SC index, however, is much more challenging due to the time-varying frequency and time-varying strength of the cardiogenic artifact, and a modification of the SC index is introduced. By warping the EEG signal, the spectrum associated with cardiogenic artifacts becomes more concentrated around the fundamental frequency and its multiples. While the EEG spectrum is also deformed due to the warping, the spectral spreading issue caused by the inevitable heart rate variability is mitigated, and hence the EEG recovery can be better evaluated. Furthermore, our algorithm also outperformed in metrics for EEG signal recovery when we compared recovered signal in the semi-real dataset to the ground truth EEG.

We further identified that the cardiogenic artifact does not only entail the strong QRS complex but also the less pronounced T-wave, which has previously been reported^9^. Thus, we highlight the importance of capturing a long enough window around the QRS complex to remove the T-wave of the cardiogenic artifact.

### Beyond artifact removal: Opportunity to capture heart related activity without ECG

Besides using brMEGA for cardiogenic artifact removal, it can also be leveraged to extract heart-rate related activity (e.g. heart rate variability measures, IHR) without the need of additional ECG. In some applications, like telemedicine and mobile health, where only limited channels are available or ECG channels around the chest might be disturbing (e.g. sport science), this extra information, when exists, might provide more health information and benefits. Even in the monitor-intense environment like an intensive care unit or operation room, this information might also be useful when other first line heart rate information is of bad quality.

### Limitations and future works

Our brMEGA approach offers several advantages in removing cardiogenic artifacts from single-channel mobile EEG data, that is a completely automatic framework to identify cardiogenic artifacts, peak detection and removal without the need of an additional ECG reference. Yet, this algorithm has clear potential to be further improved in future studies.

First, the sensitivity of the algorithm could be enhanced and validated against different approaches. Using visual identification as the ground truth of artifact existence is the step that can be improved. One problem is that strong EEG background activity might mask its presence so that the ground truth is not correct, which will mislead the SVM model and downgrade the overall performance. Therefore, while the algorithm could in theory outperform the visual capacity of the experimenter, this ultimate goal cannot be achieved in this work. Even with a better annotation, yet, this would need to be tested in different frameworks, such as with many experts and definition of a consensus rule among the independently rated expert opinions.

Second, while the brMEGA removes the artifact substantially and outperforms a classical EAS approach, there is still some remaining artifact that are hard to visualize by naked eyes. To further improve the algorithm, however, will be a combined effort of optimizing all three steps of the brMEGA, since all of them contribute to the artifact removal performance of long-term recordings.

Finally, it remains to be established whether brMEGA could be used to clean EEG signals that are contaminated by other strong artifact sources that resemble spike-like patterns, e.g. epileptic seizures.

## Material and Methods

### Used datasets

#### Single-channel EEG recordings from portable device

Thirteen EEG recordings of nocturnal sleep periods from different participants have been selected for this publication. Participants (all healthy, 63-74 years of age, 4 female) were part of an ongoing clinical trial (NCT03420677) that has been approved by the provincial ethics committee and the Swiss Agency for Therapeutic Products Swissmedic. The EEG data reported here was collected with the MHSL-Sleepband, a mobile and configurable system for high-quality biosignal recordings at home^6^. Specifically, participants used this system independently at home and therefore applied single-use auto-adhesive electrodes (Neuroline 720, Ambu A/S, DK) to different electrode positions on the scalp and face to capture EEG, EOG and EMG activity. For the purpose of this article, we will only focus on the EEG, which was placed at Fpz with a reference on the right Mastoid (M2) and the ground on the left Mastoid (M1). The signal was sampled at 250 Hz.

The neuroscientist expert in our team visually examined the EEG signal (filtered at 0.5 Hz high-pass and 40 Hz low-pass for visualization purpose only) and provided a quality over non-overlapping 4 s epochs in the sense of cardiogenic artifact. For 13 whole night polysomnograms, the neuroscientist labeled randomly selected 100 4-second epochs between the first and second hour of each recording. An epoch was labeled as 0 if no clear cardiogenic artifacts could be visually identified. An epoch was labeled as 1 when a cardiogenic artifact was visible.

### Control datasets with ECG reference

The control dataset consists of 2 clinical subjects (one 32-year-old male and one 63-year-old female) being evaluated for sleep apnea by the whole night polysomnogram but diagnosed to be free of sleep apnea. The recordings were collected from the sleep center at Chang Gang Memorial Hospital (CGMH), Linkou, Taiwan. The institutional review board of the CGMH approved the study protocol. All signals were acquired by a biosignal amplifier system from Philips Alice. These recordings are part of the Taiwan Integrated Database for Intelligent Sleep (TIDIS) project, so we refer this database as TIDIS. The simultaneously recorded EEG and ECG signals were sampled at 200 Hz. For the validation purpose, we select F4M1 channel for one case and O1A2 channel for another case that have visually identifiable cardiogenic artifacts.

### Semi-real database with EEG and ECG ground truth

The semi-real dataset consists of 7 simulated over-night recordings. The original recordings were collected from the sleep center at CGMH, Linkou, Taiwan. The institutional review board of the CGMH approved the study protocol. All signals were acquired by a biosignal amplifier system from Philips Alice, which are part of the TIDIS project as well. We randomly select 7 EEG recordings without cardiogenic artifacts and generate the semi-real recording in the following way. For the EEG, there are in total 7,146 epochs, and 28 epochs are removed by the following criterion. If there exists an interval longer than 4 seconds with the EEG magnitude less than or equal to 1.5 microvolt, that epoch is removed. We preprocess the raw ECG signal by resample it to 250 Hz and then applying a median filter with the window length 0.2 s followed by a low pass filter with the cutoff at 80 Hz. Then, select 20 seconds intervals with bSQI^32^ greater than or equal to 0.95, and we also remove the 20 seconds intervals if the maximum value of the difference between the background power of that interval and the median height of all peaks in this interval is greater than 200 microvolt or if the maximum of the difference between any peak and the median height of all peaks in this interval is greater than 120 microvolt. We concatenate them to form the clean ECG signal. If two consecutive beats have RR interval less than 0.4 s, the first one is removed. Next, we generate the cardiogenic artifact by randomly scaling the peaks in the following way. Let *R*_*i*_ be the height of the *i*th R peak in the clean ECG, and *QS*_*i*_ be the interval of the QRS interval of the *i*th heartbeat, and *SQ*_*i*_ be the interval from the S wave of the *i*th QRS to the Q wave of the (*i* + 1)th QRS, which includes the *i*th T wave. We scale the ECG over *QS*_*i*_ by 60*h*_*i*_/*R*_*i*_, where *h*_*i*_ is independently and identically sampled uniformly from [0.8, 1], and let scale the ECG over *SQ*_*i*_ by 12*h*_*i*_/*R*_*i*_. We consider this scaling since the T wave is not always there and not strong even it exists in the cardiogenic artifact contaminated EEG signal. Finally, sum the simulated cardiogenic artifact and the cardiogenic artifact free EEG signal to form the final semi-real EEG signal.

### Cardiogenic artifact detection and removal algorithm

The algorithm is composed of two main steps. The first step is determining if an epoch is contaminated by cardiogenic artifacts. The second step is detecting the location of each cardiogenic artifact and removing the cardiogenic artifact. The algorithm we consider is a modification of the shape adaptive nonlocal artifact removal (SANAR) designed for removing the stimulation artifacts during the direct cortical stimulation^23^. The main difference is that the periods between two consecutive stimulation artifacts are usually fixed and their times are known, while the periods between consecutive cardiogenic artifacts are time-varying and their times are unknown. Below we detail the algorithm step by step. Denote the digitalized EEG signal as *x*_0_ ∈ ℝ^*n*^, where we pad the recorded signal by 0 so that the overall length of the EEG signal is of the multiple of 4 seconds. We assume that the signal is uniformly sampled every Δ_*t*_> 0 second. Without loss of generality, we assume that 1/Δ_*t*_ is an integer to simplify the notation and suppose there are *L* > 0 4-second non-overlapping epochs; that is *n* = *L* × 4/Δ_*t*_.

### Step 0: Preprocessing of EEG data for algorithm

The mean of the EEG signal is removed and an IIR notch filter applied at 50 Hz (or 60 Hz, depending on where the signal is recorded) to suppress the power line interference. Then, we remove the wandering baseline by applying the median filter with a window length of 100ms, followed by a window smoothing of length 150ms. Fix Δ_*t*_> 0. If the sampling rate is different from 1/Δ_*t*_Hz, we resample the signal to 1/Δ_*t*_ Hz to balance between the computational time and a better recovery of the cardiogenic artifact information^35^. In practice, if the sampling rate is higher than 1/Δ_*t*_ Hz, the signal is downsampled to 1/Δ_*t*_ Hz, and the reconstructed cardiogenic artifact is upsampled back to the original sampling rate and subtracted from the EEG signal. We comment that if computational time is not an issue, the following algorithm can be applied to the original signal with parameters scaled accordingly. We denote the resulting signal as *x*_1_ ∈ ℝ^*n*^.

### Step 1

In the first step, we automatically determine if a given epoch is contaminated by cardiogenic artifacts or not. We first correct the polarity of the signal. To do this, we need to locate peaks in *x*_1_. We rectify the signal and take the square root of it. Denote the resulting signal as *y*_1_ and detect peaks in *y*_1_. The peak locations of *x*_1_ are determined by the peak locations of *y*_1_. If the mean value of peak heights in *x*_1_ is negative, we correct the polarity by flipping the sign of *x*_1_, so that the peaks in *x*_1_ are positive. After the correction, to strengthen the power of peaks relative to the background brainwave, we set the negative part of *x*_1_ to be zero and keep the positive part of *x*_1_. Denote the resulting signal *x*. Then we run the nonlinear-type time-frequency (TF) analysis tools, synchrosqueezing transform (SST), on *x*. We focus on the frequency band [0, *U*), where *U* ≤ 1/2Δ_*t*_. Note that 1/2Δ_*t*_ is the Nyquist rate. Suppose the frequency axis is equally spaced into *m* > 0 bins. Set Δ_*ξ*_: = *U*/*m*. The resulting time-frequency representation (TFR) by SST is denoted as *S*_*x*_ ∈ ℂ^*n*×*m*^ respectively. We then divide *S*_*x*_ into *L* submatrices associated with *L* non-overlapping epochs, denoted as 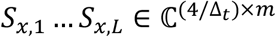.

According to the well-established theory, the cardiogenic artifact in EEG will be represented as a dominant horizontal curve in the TFR^28^. See Figure 1, right panel, for an example of the TFR of an EEG segment contaminated by cardiogenic artifacts by SST. It is clear that when the cardiogenic artifact is strong, we can easily identify a dominant curve. Thus, since the cardiogenic artifact reflects the heart beats, that curve represents the IHR, and we obtain the IHR of the subject if we can extract that curve. Since we focus on subjects without severe arrhythmia like ventricular fibrillation, we follow the normal physiological constraint and assume that the heart beats at the frequency less than 4Hz and higher than 1/3Hz; that is, the consecutive two beats are separated by at least 250 ms and no longer than 3 seconds^29^. Under this constraint, in this work, we apply the following curve extraction algorithm to determine the IHR.

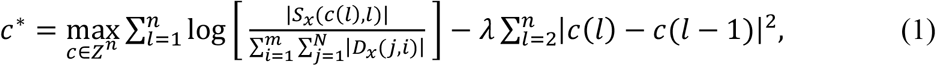

where *λ* > 0 is the penalty term chosen by the user, *Z* = {⌊1/3Δ_*ξ*_⌋ + 1, ⌊1/3Δ_*ξ*_⌋ + 2, … ,4/Δ_*ξ*_}, ⌊1/3Δ_*ξ*_⌋ means the largest integer smaller than 1/3Δ_*ξ*_, and *c*\* is a *n* dim vector representing a curve on the TFR. The first term in the function in (1) essentially fits the TFR with a curve that captures the frequency with the highest intensity at each time, while the second term imposes a smoothness constraint on the extracted curve. As a result, when cardiogenic artifacts exist, the IHR is captured, and at time *i*Δ_*t*_-th second the IHR is *c*\*(*i*)Δ_*ξ*_ Hz. When there is no cardiogenic artifact, the extracted curve represents the instantaneous frequency associated with the spectral distribution of the EEG as a random process.

With the estimated *c*\*, we apply the peak tracing algorithm^36^ to determine the location and heights of peaks from *x*. Specifically, we consider the following optimization:

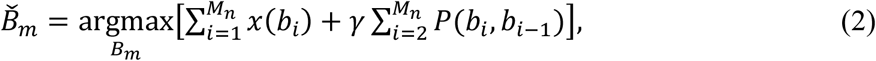

where 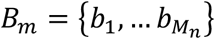 is the set of estimated peak locations, *γ* > 0 is the penalty term, and

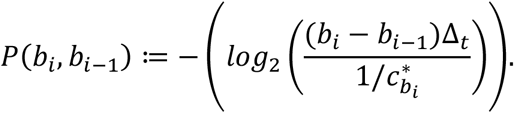

Note that *P*(*b*_*i*_, *b*_*i*−1_) is the term constraining the estimated peak locations so that they follow the estimated curve 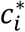. Indeed, (*b*_*i*_ − *b*_*i*−1_)Δ_*t*_ is the period between the estimated (*i* − 1)th and *i*th peaks, and 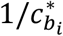 is the estimated instantaneous frequency at the estimated *i*th peak. Note that when cardiogenic artifacts exist, 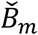 contains the locations of cardiogenic artifacts. Otherwise, we obtain random peaks associated with the EEG signal. Since we do not consider patients with severe arrhythmia, if two detected peaks are separated less than 250ms, we keep the beat with the larger amplitude and remove the other one.

Motivated by the physiological meaning of *c*\*, we design features and construct an automatic cardiogenic artifact detection algorithm from *x* and its SST. Overall, we consider the following four features associated with the cardiogenic artifact in each 4-second epoch, and then train a support vector machine (SVM) model^37^.

Since the cardiogenic artifact represents itself as a spike with the spike height generally higher than the EEG signal, the first feature we consider is the *peak height* of cardiogenic artifact. See Figure 1 for an example of the cardiogenic artifacts. For the *l*-th 4-second epoch, we take the median height of all detected peaks in that epoch to be the feature associated with the epoch. We denote this first feature as *X*_*l*_(1). This feature is obvious from the time domain.

Next, for the *l*-th epoch, we consider the optimal transport (OT) distance, or the earth mover distance to capture the strength of the cardiogenic artifacts from the TF domain as the second feature. To this end, we first prepare some quantities associated with the cardiogenic artifact in the TF domain and take the extracted curve *c*^(1)^. As is mentioned above, it is associated with the fundamental frequency of the cardiogenic artifact. We apply (1) on |*S*_*x*_|, after masking the extracted curve *c*^(1)^ by 0, to extract the instantaneous frequency of the first harmonic of the cardiogenic artifact. Denote it as *c*^(2)^ ∈ ℝ^*n*^. By iterating this process, we obtain the second and third harmonics of the cardiogenic artifact, denoted as *c*^(3)^ ∈ ℝ^*n*^ and *c*^(4)^ ∈ ℝ^*n*^. It is critical to note that the instantaneous frequencies of “harmonics” might not exactly be integer multiples of the fundamental frequency. A critical observation of cardiogenic artifact is that while cardiogenic artifacts look similar, their patterns might change from one to the other. This time-varying pattern has been extensively discussed in the literature^30^. With this preparation, we restrict *c*^(1)^ … *c*^(4)^ on the *l*-th epoch, and denote them as 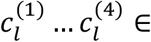 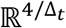. For the *k*-th sampling point of the *l*-th epoch, define the *ideal* time-varying spectrum as a *m* -dim vector *ν*_*k*_, so that 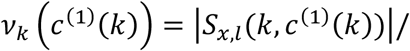 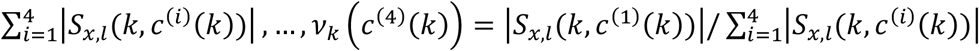 and other entries 0. We also define the estimated time-varying spectrum associated with the *k*-th sampling point as 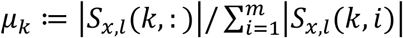. Clearly, *ν*_*k*_ and *μ*_*k*_ are both probability measures. Note that when the cardiogenic artifact is strong, the energy will be concentrated around the instantaneous frequencies of the fundamental and harmonic components. Thus, the estimated time-varying spectrum will be close to the ideal time-varying spectrum. Otherwise, they will be different. To quantify this difference at the *k*-th sampling point, we apply the OT distance^38^, which is defined as 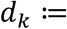 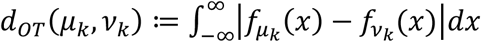, where 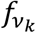 and 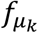 are the cumulative functions of *ν*_*k*_ and *μ*_*k*_. Thus, the first feature in the TF domain for the *l*-th epoch, *X*_*l*_(2), is defined as the median of 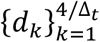. The second feature in the TF domain for the *l*-th epoch is defined similarly, but we only focus on the fundamental component; that is, for the *k*-th sampling point of the *l*-th epoch, consider a *m*-dim vector 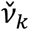, so that 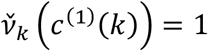 and other entries 0, and define the second feature in the TF domain for the*l*-th epoch, denoted as *X*_*l*_(3), as the median of 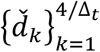, where 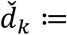 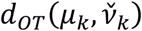. The fourth feature is the total energy over the *l*-th epoch, which is denoted as *X*_*l*_(4).

With the above four features for the *l*-th epoch, denoted as a 4-dim vector *X*_*l*_, we apply the kernel SVM with a radial basis function kernel to train an automatic cardiogenic artifact annotation system. The features are standardized by the z-score. For the reproducibility purpose, we use the SVM implementation commonly applied in practice^39^.

### Step 2

Once we confirm that the EEG signal is contaminated by the cardiogenic artifact, we apply the nonlocal Euclidean median (NLEM) algorithm to remove the cardiogenic artifact. The NLEM approach belongs to the category of template subtraction (TS), which is particularly useful when we have only single or few channels and the adaptive filter technical and blind source separation tools cannot be applied. It is designed by a critical observation that while the cardiogenic artifacts follow a similar pattern, it does vary from beat to beat. This observation motivated us to introduce the *wave-shape manifold* model^30^, which parametrizes all possible patterns of cardiogenic artifacts. By finding all cardiogenic artifacts with a similar pattern, the NLEM helps recover the cardiogenic artifact pattern more efficiently compared with the traditional TS algorithms.

Suppose there are *M* epochs that are labeled with cardiogenic artifact contamination. Assume the cardiogenic artifacts happen at *t*_1_ < *t*_2_ … < *t*_*N*_ determined by the peak tracing algorithm^36^. According to the assumption and the peak tracking algorithm, *t*_*i*_ − *t*_*i*−1_ is sufficiently large so that two cardiogenic artifacts are separated by 250ms. Before applying the OS algorithm, we apply the 5^th^ order butterworth highpass filter with the cutoff at 5Hz to the EEG signal to suppress the background EEG signal. Denote the resulting EEG signal 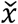. For each *t*_*i*_, determine a segment that contains the *i*-th cardiogenic artifact by 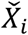 to be 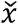 over the period [*t*_*i*_ − *L*_*L*_, *t*_*i*_ + *L*_*R*_], where *L*_*L*_ = ⌈0.2/Δ_*t*_⌉ and *L*_*R*_ = ⌈0.5/Δ_*t*_⌉. Then, form a *p* by *N* data matrix 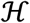, where *p* = *L*_*R*_ + *L*_*L*_ + 1 is the length of each segment and the *i*th column is the *i*-th artifact cycle.

Suppose we have 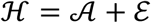, where the *i* th column of 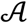 and 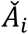 is 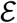 and 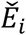 respectively, where 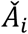 is the cardiogenic artifact and 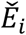 is the EEG signal. Note that after the 5Hz highpass filter, the EEG signal 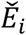 has been suppressed while 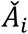 is kept less perturbed. Then, perform the optimal shrinkage (OS) algorithm with the Frobenius norm^40^ on 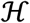. The OS result of 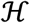, denoted as 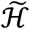 is the optimal approximation of 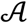 in the sense of the Frobenius norm. Thus, we can view the *i*th column of 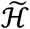 as a good estimator of the *i*th cardiogenic artifact. We can then define the following “metric” to find those EEG segments that have the same or similar cardiogenic artifacts: *d*(*X*_*i*_, *X*_*j*_) is defined as the diffusion distance^41^ between 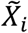 and 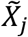, where 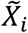 is the *i*th column of 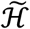.

Finally, we can apply the NLEM. For the *i*th EEG segment, denoted as *X*_*i*_, find all EEG segments with similar cardiogenic artifacts by finding the *K* EEG segments with the smallest *d*(*X*_*i*_, *X*_*j*_). Denote those *K* EEG segments as 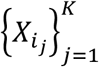. Then, find the median of 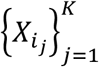, denoted as 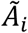, as the final recovered cardiogenic artifact. Finally, we can recover the EEG signal by removing 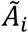 from the original EEG signal, with the common stitching policy^23^.

#### Comment

The key step for the NLEM is finding all EEG segments that have similar cardiogenic artifacts. However, it is not an easy job due to the existence of the EEG signal. To appreciate this, for the EEG signal *x* over the period [*t*_*i*_ − *L*_*L*_, *t*_*i*_ + *L*_*R*_] denoted as *X*_*i*_, we write it as the sum of two parts, that is, *X*_*i*_ = *A*_*i*_ + *E*_*i*_, where *A*_*i*_ is deterministic that models the cardiogenic artifact and *E*_*i*_ is stochastic with mean 0 that models the EEG signal with possibly the inevitable noise. The usual *L*^2^ distance between two EEG segments, *X*_*i*_ and *X*_*j*_, is 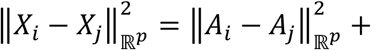 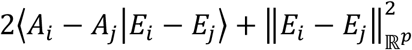. Clearly, if two cardiogenic artifacts are close, 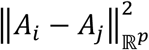 will be close to 0 and ⟨*A*_*i*_ − *A*_*j*_|*E*_*i*_ − *E*_*j*_⟩ will be a random variable with mean 0 and small variance. However, the noise term 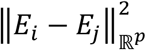 will be nonzero and might deviate the neighboring information determined by 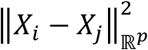. This domination might render two EEG segments with similar cardiogenic artifact patterns to have a huge distance. As a result, the neighbors determined by the usual *L*^2^ distance might not have similar cardiogenic artifact patterns. Thus, we need a new method to find those EEG segments with the same or similar cardiogenic artifacts. To this end, we propose to apply the currently developed OS algorithm^40^ to suppress *E*_*i*_ from *X*_*i*_. OS can be viewed as a nonlinear matrix denoising algorithm that removes the noise in an efficient way. Note that in the proposed algorithm, we also applied a 5Hz highpass filter before applying OS to help suppress the impact of EEG signals.

### Quantification indices

The AR index for the *i*th cardiogenic artifact, denoted as *AR*_*i*_, is defined as

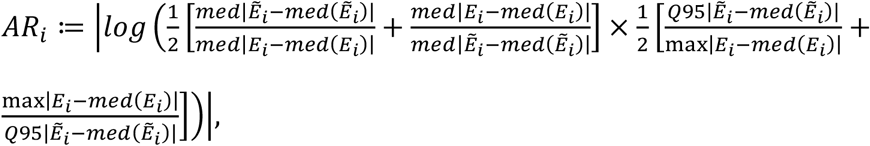

where 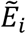 is the estimated EEG signal recovered over the *i*-th cardiogenic artifact, *E*_*i*_ is the concatenation of true EEG signal over the interval *without* any cardiogenic artifact from *i* − 5, … , *i* + 5 cardiogenic artifacts, med means median, Q95 means the 95% quartile, and max means the maximal value. As is indicated above, the cardiogenic artifact might have a “T” wave, thus, depending on the heart rate, sometimes the true EEG signal might not exist over an artifact to artifact interval. The AR index of a good reconstruction algorithm should be close to 0.

The SC index measures the performance of the algorithm in the frequency domain. The original SC index for the EEG signal is defined as the ratio of the power change, relative to raw signal, over the fundamental frequency and the harmonics of the stimulation frequency to the energy over the rest of the frequencies in the band (1-100 Hz). While this definition works well for stimulation artifacts removal problem, however, it is not suitable for cardiogenic artifact, due to spectral spreading caused by the existence of heart rate variability. To avoid this issue, we propose the following modified SC (mSC) index. First, the EEG signal is warped in the time domain by the estimated instantaneous heart rate (IHR) so that the cardiogenic artifacts have a constant frequency that is equivalent to the mean of the estimated IHR. Second, the SC index is evaluated from the warped EEG signal. The EEG signal and warped EEG signal are both sampled at 250 Hz. The mSC index is defined as the ratio of the power change, relative to raw signal, over the fundamental frequency and the harmonics of the stimulation frequency to the energy over the rest of the frequencies in the band (1-100 Hz). The band 1-100 Hz is chosen since it is sufficiently wide to cover spectrum of interest, including delta (1-4 Hz), theta (4-8 Hz), alpha (8-13 Hz), beta (13-30 Hz) and gamma (30-50 Hz), even if the spectrum of warped EEG signal is clearly different from that of the original EEG. The SC index should be small if the reconstruction quality is good.

To apply the AR and SC indices, we adhere to the following protocol. For each 20 s epoch, we first get 5 prediction results for all non-overlapping 4 sec sub-epochs. In each 20-sec epoch, if more than and equal to 3 sub-epochs are determined to have cardiogenic artifacts, then we apply the cardiogenic removal algorithm to that 20-sec epoch and evaluate the AR and SC indices. For the AR index, it is evaluated for each peak in the 20 sec epoch and take the median of them to represent the AR index for that 20-sec epoch. For the SC index, we calculate the index over the 20-sec epoch.

The above AR and SC indices are designed to evaluate the algorithm performance when the ground truth EEG signal is not available. When the true EEG signal is available, we apply the following four metrics^33^ to quantify how well the EEG is recovered from the cardiogenic artifact contaminated EEG signal. Denote the true EEG as a ∈ ℝ^N^ and the estimated EEG as 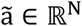. Here, the estimated EEG can be the one after applying brMEGA or EAS, or without applying any algorithm. The first index is the normalized mean square error (NMSE), which is defined as

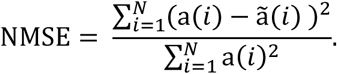

Clearly, the smaller NMSE is, the better the EEG is estimated by 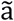. The second index is the cross correlation between a and 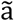 , denote as ρ. The better 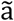 estimates a, the closer ρ is to 1. The third and fourth indices are *signal-to-noise ratio* (SNR) and *peak-signal-to-noise ratio* (PSNR), which are defined by

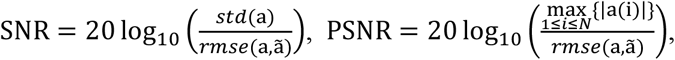

where 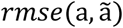 is the root mean square error between a and 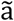. Clearly, the higher the SNR and PSNR, the better the recovery is.

To report NMSE, we divide the signal into non-overlapping 20 second epochs, and we take the median value of all NMSEs in each epoch, which represents the algorithm performance over that epoch. The same procedure is applied to ρ, SNR and PSNR in the same way. In the end we have four indices to quantify the cardiogenic artifact, and hence EEG, recovery quality over each epoch.

## Supplementary results

To further evaluate the proposed automatic determination of existence of cardiogenic artifacts, the leave-one-subject-out cross validation was carried out. In the leave-one-subject-out cross validation, the validation set and training set are determined on the subject level; that is, the training set and the validation set contain different subjects. The result is shown in Supplementary Table 1, where the sensitivity is 0.93, the specificity 0.69, and the F1 score is 0.84 and the overall accuracy is 0.8. Compared with the 5-fold cross validation shown in Table 1, the result is compatible and encouraging. Note that while the leave-one-subject-out cross validation scheme is challenged by the inter-individual variability, this scheme is close to the real-world scenario, particularly when there is no available information for a newly arrival subject.

### Implementation details and parameters

The proposed brMEGA algorithm is implemented in Matlab, and we adhere to the following parameters in this work. We set 1/Δ_*t*_= 250. In the nonlinear-type TF analyses, including SST, the window is Gaussian and the Full width at half maximum is set to be 10s, and the maximal frequency is set to be *U* = 25 Hz. We fix the frequency resolution at Δ_*ξ*_: = *U*/*m* = 0.02 Hz. Set *λ*=1 in the curve extraction algorithm (1), We set the parameter γ = 20 in the peak tracing algorithm in (2). For the kernel SVM classifier, we apply the following parameters: the cost is set to 1, Gamma is set to 0.2, and the weights are set to be 1 and 3 for the cardiogenic artifact free epochs and epochs with cardiogenic artifacts respectively. We do not carry out any grid optimization search for the parameters to avoid over-fitting. In the NLEM, we take *K* = 100.

To avoid the possible interaction between the EEG signal and the sleep dynamics and to speed up the computation, we suggest to randomly permute the cardiogenic artifacts and then perform OS on every non-overlapping 500 cardiogenic artifacts. To further speed up the computation, the NLEM step is suggested to carried out in the following way. We first divide the EEG segments 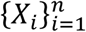 into non-overlapping groups containing consecutive 250 EEG segments. Suppose *X*_*i*_ is in the *l*th group. Find *K* EEG segments with similar cardiogenic artifacts in the sense of the smallest diffusion distance from the *l*th and the (*l*+1)th groups.

The power spectral density is calculated using Welch’s method, featuring 5000 discrete Fourier transform points, Hamming windows of 5000 samples, and 50% overlapping. The 5000 point Fourier transform is chosen to reflect a 5 second window for a frequency resolution of 0.2 Hz. Spectrogram plots (Figure 5) were derived using a multitaper approach to optimally illustrate the nocturnal neurophysiological EEG dynamics as well as the artifact over the whole recording period^42^. Specifically, we used the *mtspecgramc* function from the Chronux open source library (http://www.chronux.org/). As previously optimized for whole night spectral representation, we chose parameters of 30 s windows spaced at 5-s intervals, spectral resolution of 1 Hz, the time-half-bandwidth product of 15, and the number of tapers of 29^42^.

For the algorithm performance evaluation, the mean and standard deviation of the AR and SC indices for different algorithms were reported. For each subject, the null-hypothesis that one algorithm does not perform better than the other one in the sense of AR or SC index is tested by the paired-sample t-test. We view p value < 0.05 as statistical significance. To compare recovery of the EEG in the semi-real dataset, we first evaluated whether EAS and brMEGA improved EEG signal quality compared to the uncleaned EEG. Therefore, we performed a linear mixed model using method as a fixed factor (levels: uncleaned, brMEGA, EAS) and subject and epoch as a nested random factor. To verify whether the cleaning with brMEGA was different to EAS we took the difference of each epoch relative to the uncleaned signal and performed a linear mixed model of these differences with the fixed factor cleaning method (levels: brMEGA, EAS) condition and subject as a random slope factor.

## Data availability

The datasets analyzed during the current study are available from the corresponding author on reasonable request. In the future, the dataset will be made available in a repository.

## Code availability

The code of brMEGA is currently available via request but will be made open source in the future.

## Acknowledgements

This work was conducted as part of the SleepLoop Flagship of Hochschulmedizin Zürich funded in part by the Schweizerische Hirn Stiftung and the Swiss National Science Foundation (P3P3PA_171525 and PZ00P3_179795 to CL). The authors thank all the participants for taking part in the study and R. Büchi, N. Demarmels, G. Hoppeler, J. Kurz, and E. Silberschmidt for performing the recruitment and data collection. SleepLoop consortium members provided uncountable discussions and feedback. This work and the Taiwan Integrated Database for Intelligent Sleep (TIDIS) project are partially supported by Ministry of Science and Technology 109-2119-M-002-014-, NCTS Taiwan. Neng-Tai Chiu is partially supported by the Undergraduate Summer Research Program hosted by National Center for Theoretical Sciences, Mathematics Division, Taiwan.

