## Appendix for "Get rid of the beat in mobile EEG applications: A framework towards automated cardiogenic artifact detection and removal in single-channel EEG"

**Table A1**

| | **Visual: artifact (1) (576)** | **Visual: no artifact (0) (724)** | |
|---|---|---|---|
| **Prediction: artifact (1)** | 542 | 214 | Precision = 0.72 |
| **Prediction: no artifact (0)** | 34 | 510 | False predictive value = 0.94 |
| | Sensitivity = 0.95 | Specificity = 0.70 | Accuracy = 0.81<br>F1 score = 0.84 |

**Table A1:** The overall confusion matrix of the 5-fold cross validation.

**Figures A1-A12**

**Combined Figure legend A1-A12:** TF representation of the original EEG signal (all-night nocturnal recording, left) and after removing the cardiogenic artifact (right) for 12 participants.

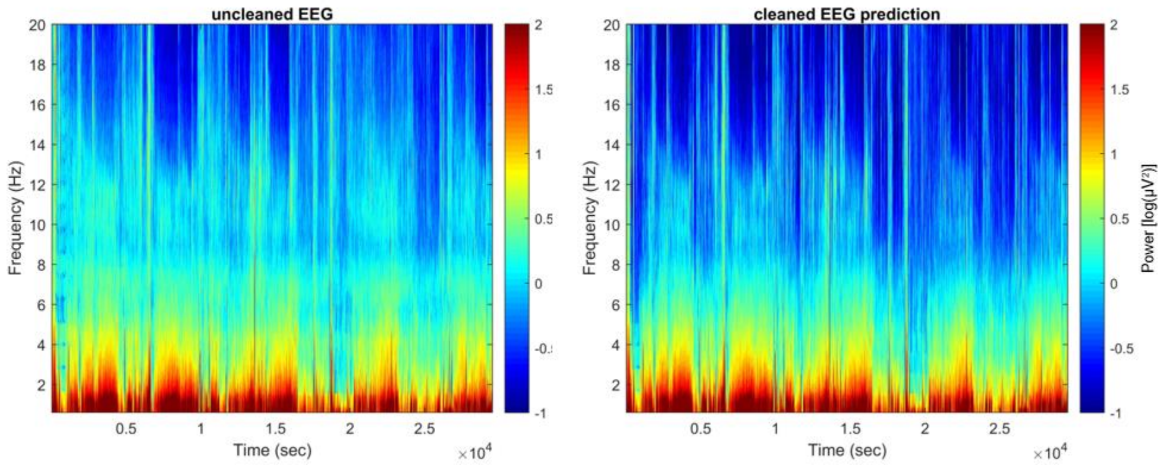

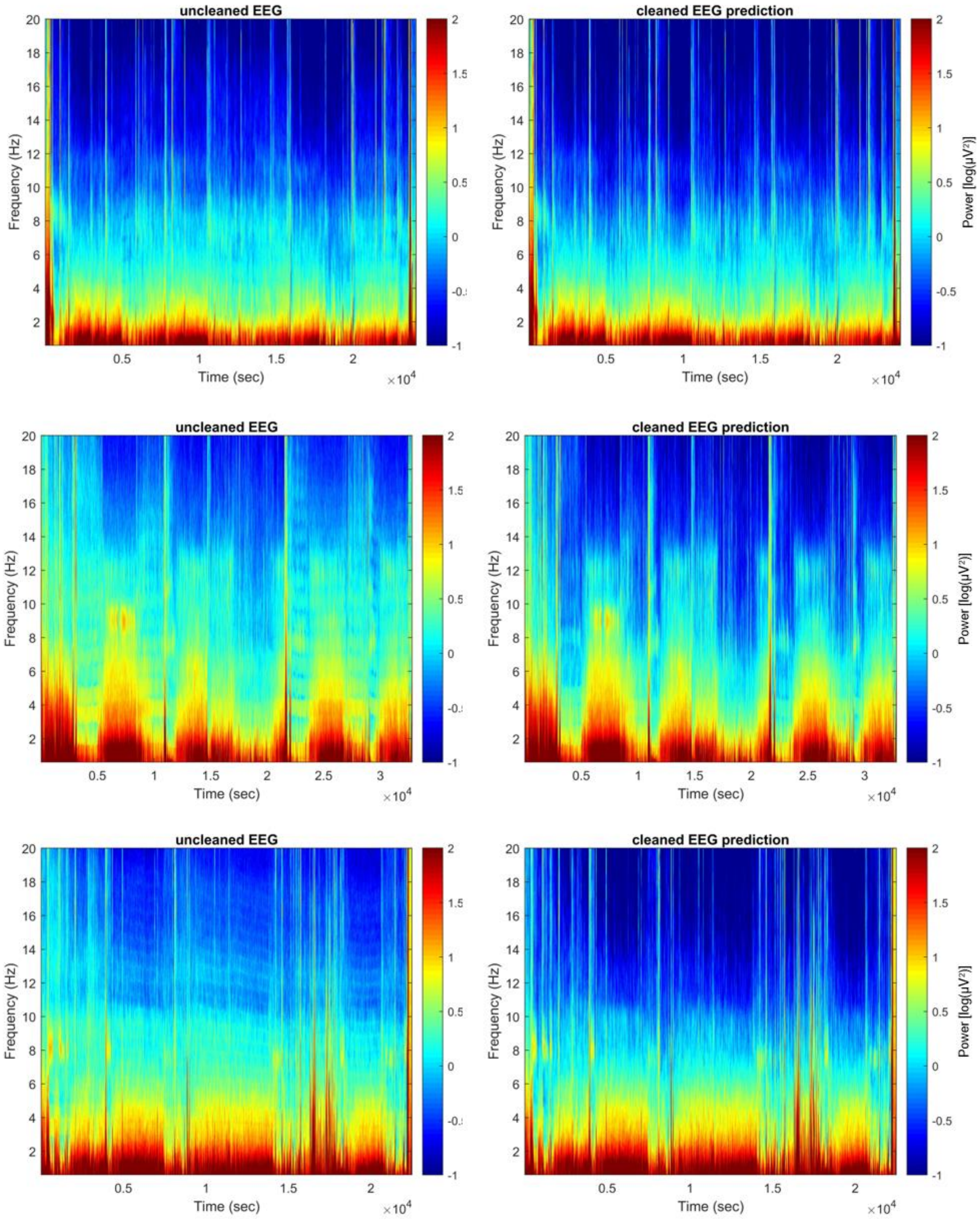
uncleaned EEG
cleaned EEG prediction
uncleaned EEG
cleaned EEG prediction
uncleaned EEG
cleaned EEG prediction
Frequency (Hz)
Time (sec)
×10^4
Power [log(μV²)]

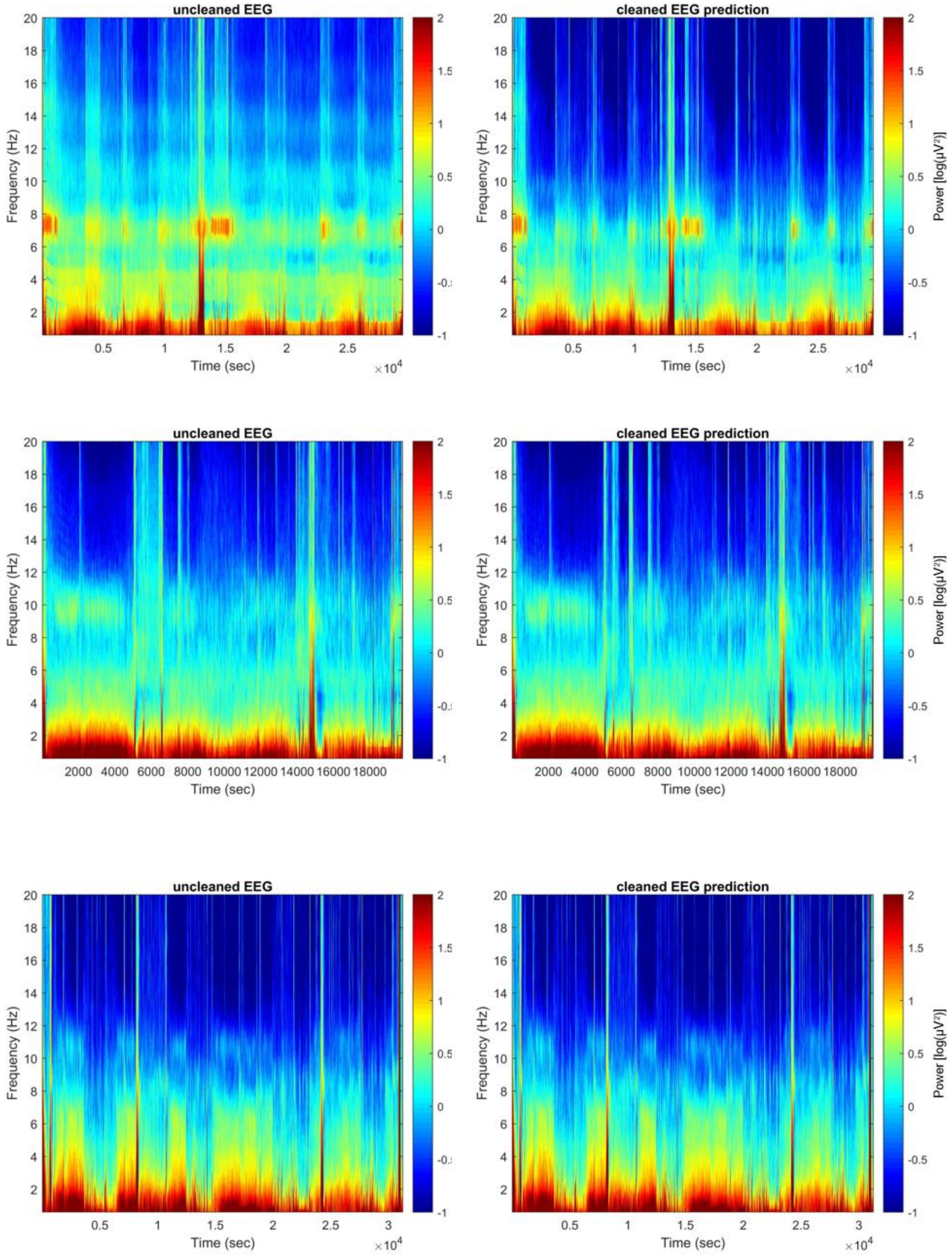
uncleaned EEG
cleaned EEG prediction
Frequency (Hz)
Time (sec)
×10^4
Power [log(μV²)]

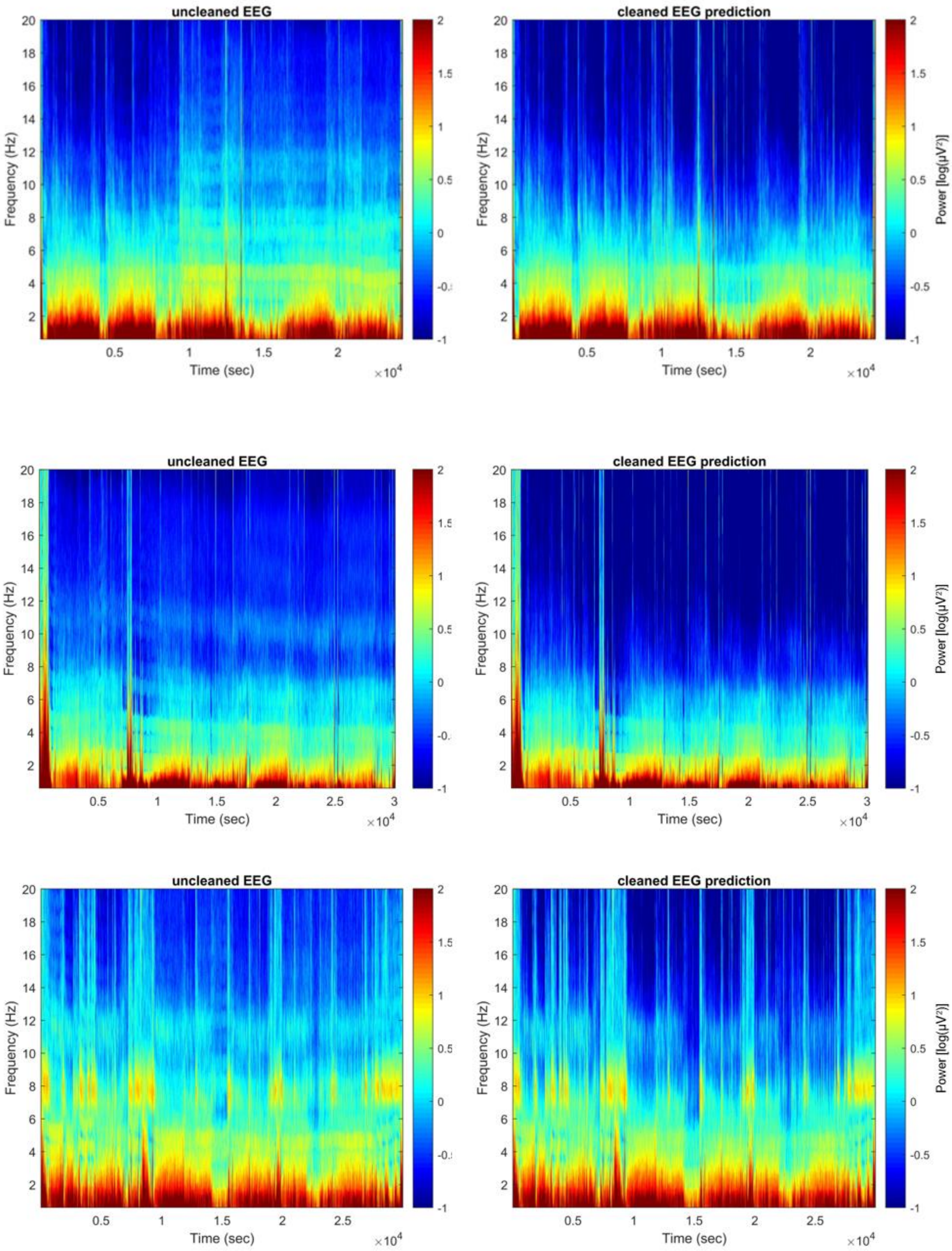
uncleaned EEG
cleaned EEG prediction
uncleaned EEG
cleaned EEG prediction
uncleaned EEG
cleaned EEG prediction
Frequency (Hz)
Time (sec)
×10^4
Power [log(μV²)]

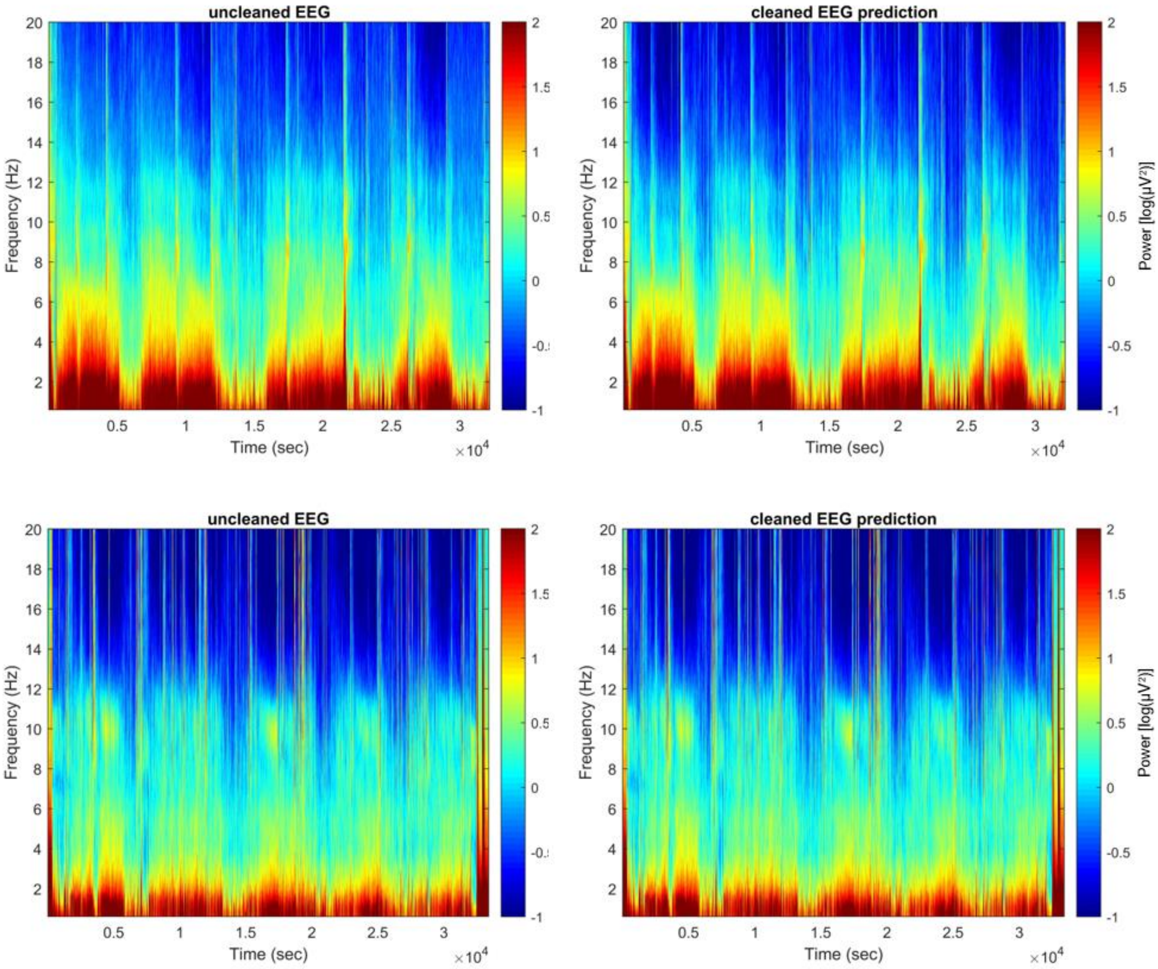
uncleaned EEG
cleaned EEG prediction
uncleaned EEG
cleaned EEG prediction
Frequency (Hz)
Time (sec)
×10^4
Power [log(μV²)]
20
18
16
14
12
10
8
6
4
2
0.5
1
1.5
2
2.5
3
-1
-0.5
0
0.5
1
1.5
2

**Figure A13**

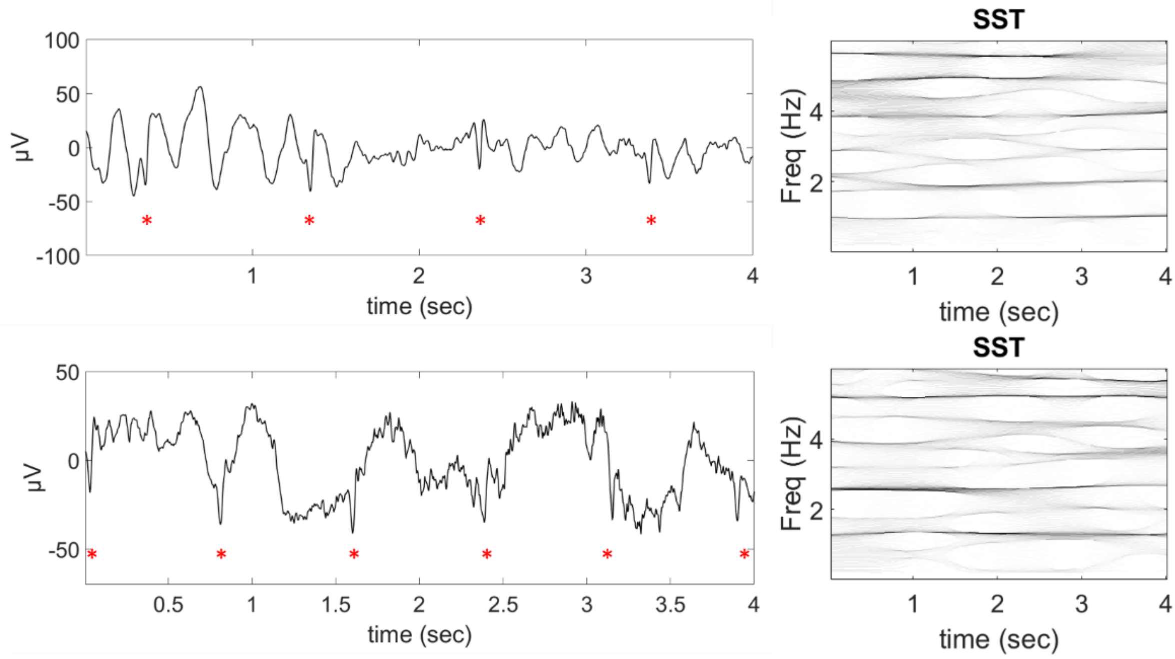


**Figure A13:** Two exemplary 4s epochs (left: temporal representation and right: SST representation) from two participants that were rated to not have a clear artifact by the neuroscientist but were predicted by the algorithm to have. Even though not clearly visible due to strong EEG activity, rhythmic peaks (marked with red Asterix) are observable that likely capture small ECG artifacts.
